# Nutrient composition and bioaccumulation of an edible aquatic insect, *Pantala* sp. (Odonata: Libellulidae) from the rice field

**DOI:** 10.1101/2021.12.26.474203

**Authors:** Witwisitpong Maneechan, Taeng On Prommi

## Abstract

Numerous edible aquatic insects have unanticipated nutraceutical potential and are consumed in a variety of Thai locations. The proximate composition, amino acid, fatty acid, mineral, and heavy metal content of *Pantala* sp. (Odonata: Libellulidae) aquatic edible nymphs were determined using standard analytical methods in this study. *Pantala* sp. had a proximate protein content of 445.14±0.04 %, a fat content of 4.93±0.05 %, an ash content of 5.24±0.03 %, a moisture content of 35.11±0.09 %, and a total carbohydrate content of 9.60±0.11 %. Total energy was 263.25±0.20 kcal/100 g, with fat energy accounting for 44.37±0.43 kcal/100 g. Inductively Coupled Plasma-Optical Emission Spectroscopy (ICP-OES) analysis revealed that this insect was high in phosphorus, iron, and copper for human consumption. In comparison to other edible insects studied, they were also excellent calcium sources. Agilent 7890B Gas Chromatograph (GC) analysis revealed that it contains 236.67 mg/100g of omega-3 and 523.32 mg/100g of omega-6. While the amino acids examined using High Performance liquid Chromatography contained all essential amino acids. ICP-OES was used to determine the levels of cadmium (Cd), lead (Pb), and arsenic (As). *Pantala* sp. had the highest concentration of As (average = 2.827 ± 0.289 mg kg_-1_), followed by Cd (0.164 ± 0.007 mg kg_-1_) and Pb (0.158 ± 0.015 mg kg_-1_). Although the insects have nutraceutical potential, they also have toxic heavy metals in trace amounts, with the exception of As. This work could serve as a nutritional reference for local consumers interested in entomophagy.

## Introduction

According to the FAO [1], the global population is predicted to reach 9.1 billion people in 2050, up 34% from now. When combined with rising economic affluence and purchasing power, the FAO forecasts that global food production will need to increase by 60% from current levels by 2050 to meet global food demands. Entomophagy is another option for dealing with these issues [2].

Entomophagy is practiced in almost every country, including Africa, Asia, and Australia [3]. According to studies, insects are a good source of protein, fat, carbohydrates, and other nutrients [4–5]. Aquatic insects are one of the most popular types of entomophagy among consumers, owing to their taste and widespread availability [6]. Insects with aquatic larvae have been documented as human food in 48 countries throughout the world. Macadam and Stockan [7] reported that Coleoptera (79 sp.) has the most edible food insect species, followed by Odonata (58 sp.) and Hemiptera (55 sp.). In Thailand, residents in the northeast consume around 150 insect species, whereas people in the north consume about 50 insect species. The richness of edible aquatic insects in Thailand is the subject of this study, which is a consequence of the country’s wide ecological and climatic diversity. During the rainy season, these insects can be found in rice fields, forested areas, natural ponds, and streams [8–9]. True bugs, beetles, and dragonfly nymphs are among the most popular insects. According to a previous study, dragonfly larvae (Libellulidae, Aeshnidae, and Gomphidae) are consumed as food in China [3, 7, 10], India [11], Philippines [12], Laos [13–14] and Thailand [9]. Thailand is expected to have ten edible species, but only five have been identified, and only three have nutritional data accessible. Among the species of dragonfly, *Anax parthenope* (Selys) is the most common edible species. Several researchers have recorded the use of aquatic insects as food, but there are few investigations into their nutritional value and bioacumulations. Therefore, this study investigates the proximate composition, amino acids, fatty acids, minerals, and toxic heavy metal contents of commonly eaten edible aquatic insects, *Pantala* sp. (Odonata: Libellulidae), from the rice fields of central Thailand.

## Materials and methods

### Sample collection and preparation

Specimens of *Pantala* sp. (Odonata: Libellulidae) final instar nymphs (Fig 1) were captured from a rice field using an aquatic D-net before they left the water and emerged as adults. For nutritional analysis, they were identified using taxonomic keys, rinsed with clean water, and sun-dried.

**Fig 1.**
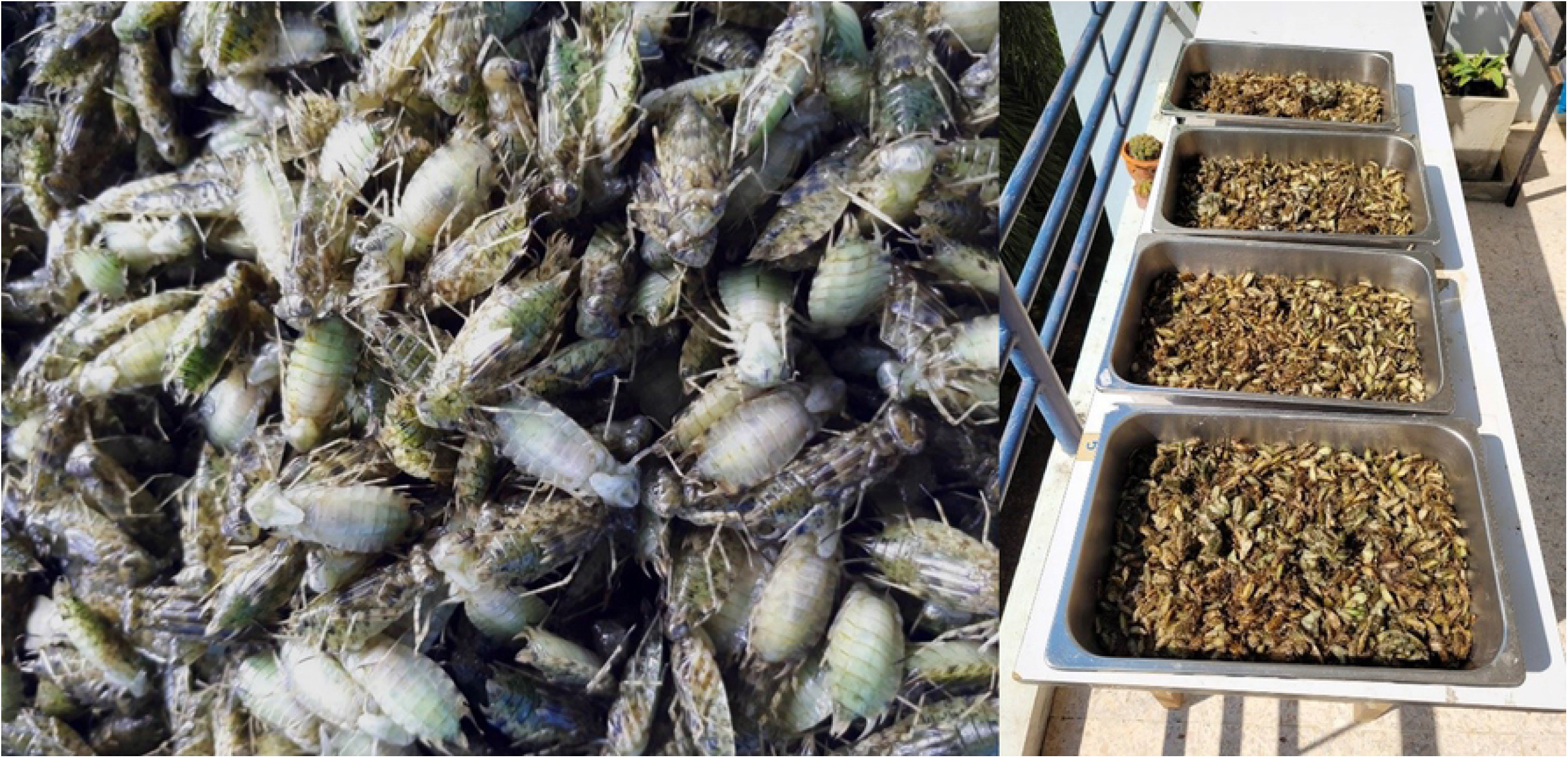
*Pantala* sp. nymph (Odonata: Libellulidae)

### Proximate analysis

The AOAC techniques 925.45, 991.20, 2003.05, and 923.03 were used to measure moisture, protein, fat, and ash content, respectively. The energy from fat in insects in Kcal/100 g was calculated by multiplying the percent fat by 9.0. [15]. The total energy of insects in kcal/100 g was calculated according to FAO [1].

### Mineral analysis

The samples were digested for 1 hour at room temperature with nitric and perchloric acid (HNO_3_: HClO_4_ = 2:1), filtered with distilled water, the final volume made up of 50 ml, and the filtrate was utilized for mineral analysis. Inductively Coupled Plasma-Optical Emission Spectroscopy (ICP-OES) was used to determine mineral and hazardous heavy metal concentrations.

### Amino acid analysis

The samples were hydrolyzed for 24 hours at 110°C with 5 mL of 6 M HCl. The amino acid composition of the hydrolyzed samples was determined using High Performance liquid Chromatography (HPLC-Agilent 1260 Infinity series, Fluorescence Detector). The HPLC’s operating conditions were as follows:

|  |  |
| --- | --- |
| Instrument | High Performance liquid Chromatography (Agilent Technologies, Germany) |
| Column | Princeton SPHER C30, 250 mm, ID 4.6 mm |
| Detector | 17 amino acid standard using FLD at Ex: 340, Em: 450; Proline using FLD at Ex: 270, Em: 316 |
| Column temperature | 30 °C |
| Flow rate | 0.7 ml/min |
| Injection volume | 16 and 10 µl |
| Moblie phase | Gradient NaH<sub>2</sub>PO<sub>4</sub>: (MeOH+ACH+H<sub>2</sub>O: 45+40+15) |
| Run time | 65 min |
| Post time | 5 min |

### Fatty acid analysis

The fatty acid profile was determined using an Agilent 7890B Gas Chromatograph (GC) and modified Compendium of Methodology for Food Analysis methods [16].

### Statistical analysis

The Statistical Package for Social Sciences (SPSS) version 19.0 was used to report the data as mean ± standard deviation.

## Results and Discussion

### Proximate compositions

The moisture, protein, total carbohydrate, lipid, ash, total energy, and energy from fat content of edible aquatic insects (*Pantala* sp.) were investigated. These results are presented in Table 1.

**Table 1.** Proximate compositions of *Pantala* sp.

| Proximate compositions | Mean±SD | Recommended Adult Reference (g/day)* |
| --- | --- | --- |
| Protein, % (factor 6.25) | 45.14±0.04 | 50 g |
| Fat, % | 4.93±0.05 | 70 g |
| Ash, % | 5.24±0.03 | - |
| Moisture, % | 35.11±0.09 | - |
| Total carbohydrate, % | 9.60±0.11 | 260 g |
| Total energy, (Kcal/ 100 g) | 263.25±0.20 | 2000 kcal/day |
| Energy from fat, (Kcal/ 100 g) | 44.37±0.43 | - |

Results are in means of triplicate determinations ± SD. \*Data derived from Williams and Williams [3].

In terms of moisture and ash content, *Pantala* sp. had a greater percentage of moisture (35.11%) and ash content (5.24%) than edible aquatic insects (Odonata: Libellulidae) reported by Shantibala *et al*. [17] and Narzari *et al*. [18]. This means that meals with a high moisture content are more susceptible to microbial growth and enzyme activity [19]. The mineral composition is determined by the ash content [6]. On a dry weight (DW) basis, the protein content of *Pantala* sp. was lower than that of *Sympetrum* sp., *Crocothemis servilia* (Drury), *Gomphus cuneatus* and *Lestes praemorsus* reported by Ying *et al*. [10], Shantibala *et al*. [17] and Narzari *et al*. [18]. Protein concentrations in aquatic insects such as Odonata can range between 40 and 65 percent [4,17]. However, the protein content of *Pantala* sp. was found to be higher than that of other edible aquatic insect species consumed in India (Coleoptera, Hemiptera, and Nepidae) [6]. Furthermore, when compared to all edible insects studied by Rahman *et al*. [20], the protein content of *Pantala* sp. was higher. Overall, the protein content of edible aquatic insects was comparable to that of chicken (42.58%), beef (35.31%), eggs (48.51%), pork (47.58%), and fish (49.63%) [21].

Lipids are essential because they provide 9 kilocalories per gram of energy [6]. The percentage of lipid content in *Pantala* sp. (4.93%) reflects its calorific value, which contributes 44.37 kcal/100 g of energy from fat. Lipids are one of the principal structural components of living cells [22]. *Pantala* sp. had a lower carbohydrate content (9.60%) than edible insects described by Narzari and Sarmah [18], but it was greater than edible insects reported by Rahman *et al*. [20] and Akullo *et al*. [23].

A total of about 263.25 kcal of energy was provided by 100 g, with approximately 1,149 nymphs. Insects are a good source of energy, with 400–500 kcal per 100 g of dry matter [24]. In comparison to the typical daily requirements of important dietary components [100 g of dried insect bodies is around 1/2 a cup], the protein yield of *Pantala* sp. is fairly high (90.28%) (Table 1). As a result, insects may be a good source of protein for those who are malnourished. Carbohydrate yield, on the other hand, is low. However, the carbohydrate amounts found in this study are higher than those found in the Shantibala *et al*. [17] report on five edible aquatic insects. The presence of carbohydrate content (9.60% [g/100g]) in *Pantala* sp. is evidence of a suitable dietary supply of carbohydrates, despite the fact that insects are not a good source of carbohydrates due to human demands of about 260 g/day [3, 25–26]. As a result, edible aquatic insects are high in macronutrients (protein, fat, and energy).

### Mineral profile

The most abundant mineral profile in *Pantala* sp. was potassium (625.26 mg/100 g), followed by sodium, phosphorus, calcium, and magnesium, respectively (Table 2). Narzari *et al*. [18] found that aquatic insects such as *Sympetrum* sp. supplement a significant amount of macronutrients such as potassium (1,690.05 mg/100g), phosphorus (626.78 mg/100g), and sodium (240.80 mg/100g). Both potassium and sodium, as well as magnesium, are electrolytes that aid in the normal functioning of nerves and muscles. These substances are located in the extracellular and intracellular fluids. Magnesium is involved in nearly all of the cell’s major metabolic and biochemical systems, including muscular contraction and relaxation, neurological function, and neurotransmitter release [27].

**Table 2.**
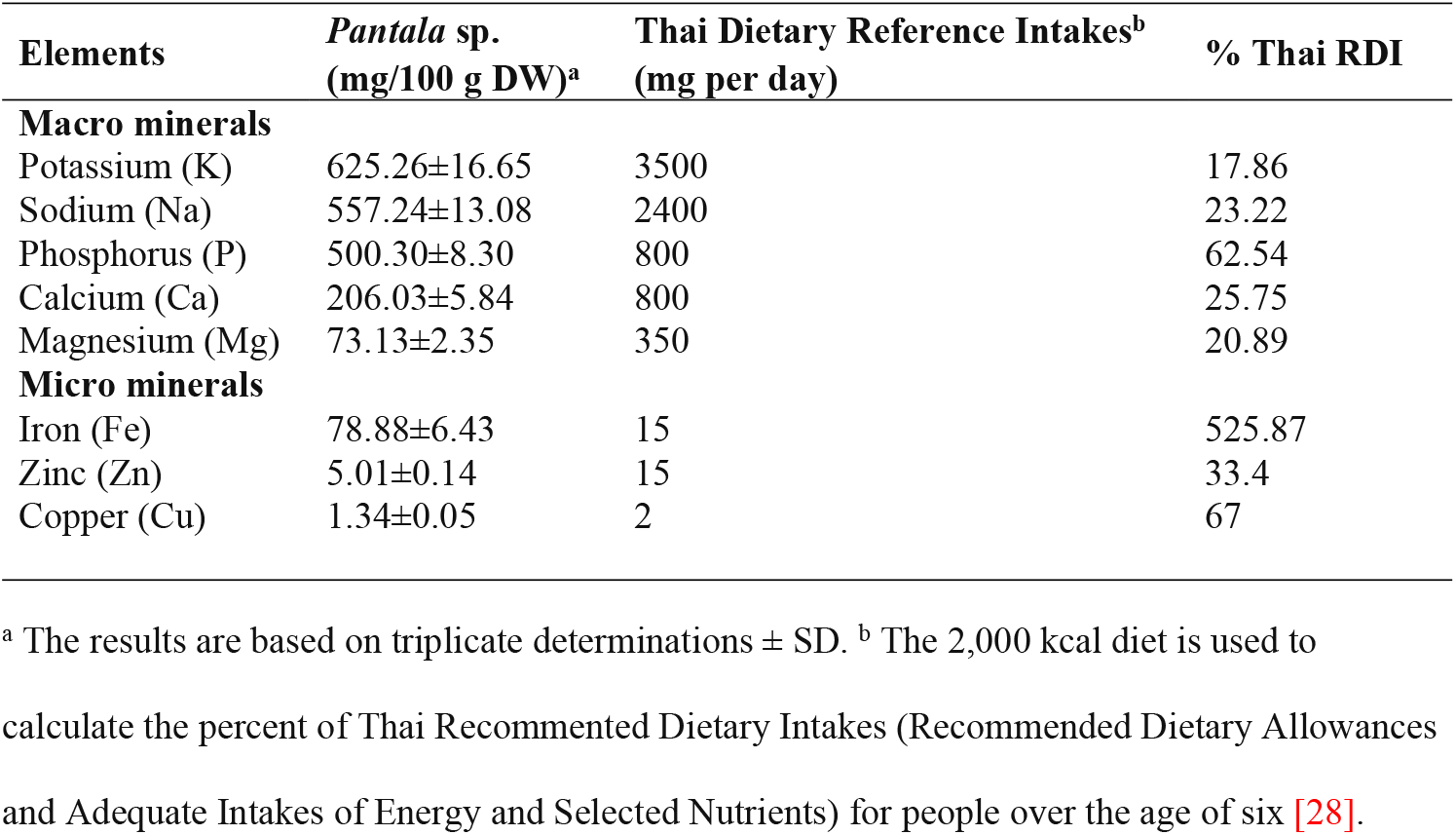
Minerals of *Pantala* sp. mg/100 g dry weight.

| Elements | *Pantala* sp.<br>(mg/100 g DW)<sup>a</sup> | Thai Dietary Reference Intakes<sup>b</sup><br>(mg per day) | % Thai RDI |
| --- | --- | --- | --- |
| **Macro minerals** |  |  |  |
| Potassium (K) | 625.26±16.65 | 3500 | 17.86 |
| Sodium (Na) | 557.24±13.08 | 2400 | 23.22 |
| Phosphorus (P) | 500.30±8.30 | 800 | 62.54 |
| Calcium (Ca) | 206.03±5.84 | 800 | 25.75 |
| Magnesium (Mg) | 73.13±2.35 | 350 | 20.89 |
| **Micro minerals** |  |  |  |
| Iron (Fe) | 78.88±6.43 | 15 | 525.87 |
| Zinc (Zn) | 5.01±0.14 | 15 | 33.4 |
| Copper (Cu) | 1.34±0.05 | 2 | 67 |

<sup>a</sup> The results are based on triplicate determinations ± SD. <sup>b</sup> The 2,000 kcal diet is used to calculate the percent of Thai Recommended Dietary Intakes (Recommended Dietary Allowances and Adequate Intakes of Energy and Selected Nutrients) for people over the age of six [28].

*Pantala* sp. had a calcium content of 206.03 mg/100g, which was higher than the calcium content of all edible insects reported by Das and Hazarika [6], Shantibala *et al*. [17], Narzari *et al*. [18], Akullo *et al*. [23], Idowu *et al*. [26], Adeduntan [29], Rumpold and Schlüter [30], Adámková et al. [31] and Das and Hazarika [32]. Calcium is the most abundant primary mineral in the human body, accounting for 1.5-2% of total body weight as bones and teeth. A calcium deficit in adults can cause osteoporosis, which is marked by bone loss and skeletal discomfort [33]. In the case of calcium insufficiency, edible aquatic insects (*Pantala* sp.) could be a potential source of calcium to supplement the diet. It is also used as a replacement for people who have a milk allergy or sensitivities to other calcium-rich foods. The recommended daily intake (RDI) amount for adults is 800-1000 mg of calcium per day [28, 34]. The source of calcium would be *Pantala* sp., which would provide between 20.60 and 25.75 percent of the daily requirements. Moreover, edible aquatic insects can be utilized as a calcium supplement for your child’s growing body, as their bones and teeth are still developing.

Phosphorus is the second most abundant major mineral in the body. It is found in every cell and part of DNA and RNA. The main role of phosphorus, together with calcium, is to aid in the calcification of bones (the skeleton contains 85% of the body’s phosphorus) and to assist the body in energy production [35].

*Pantala* sp. is high in iron, zinc, and copper, all of which are important minerals for human health. The iron content of *Pantala* sp. was 78.88 mg/100g, which was higher than those of all aquatic insects reported by Das and Hazarika [6] and Narzari *et al*. [18]. Mineral deficiencies (particularly Fe and Zn) are frequent health problems around the world, resulting in major health issues like anemia, poor pregnancy outcomes, and stunted growth, among other things [36]. Consuming 100 g of *Pantala* sp. would provide consumers with a supplement of macronutrients (potassium, sodium, phosphorus, calcium) as well as micronutrients (iron, zinc, copper) for optimal health and nutrition.

### Amino acid composition

Table 3 shows the amino acid content of *Pantala* sp. (g/100 g dry weight). Our samples contained all of the essential amino acids when compared to an aquatic edible insect, *Sympetrum* sp. (Odonata: Libellulidae) from Assam, northeast India [18]. Methionine, lysine, leucine, isoleucine, tryptophan, valine, threonine, phenylalanine, and histidine are the essential amino acids (required for the young, but not for adults). Among the essential amino acids detected, histidine is quantitatively the most abundant, with histidine being essential for the growth of infants and young children [37]. According to Igwe *et al*. [38], insect larvae are high in essential amino acids such as lysine and threonine, which are deficient in some cereals and vegetables [14]. This finding indicates that *Pantala* sp. has a high lysine content. The levels of lysine and histidine were higher than in commonly consumed meat animals such as beef, pork, and chicken. *Pantala* sp., for example, contains 2.25 g/100g lysine and 0.82g/100g histidine in beef meat, 1.80 g/100g lysine and 0.82 g/100g histidine in pork meat, and 1.79 g/100g lysine and 0.69g/100g histidine in chicken meat [39].

**Table 3.** *Pantala* sp. amino acid (AA) content (g/100 g dry weight).

| Essential AA | Total EAA | Non-Essential AA | Total NEAA |
| --- | --- | --- | --- |
| Threonine | 1.53 ± 0.13 | Aspartic acid | 2.63 ± 0.25 |
| Valine | 1.65 ± 0.12 | Glutamic acid | 4.05 ± 0.39 |
| Methionine | 0.41 ± 0.04 | Serine | 1.57 ± 0.11 |
| Isoleucine | 1.42 ± 0.10 | Glycine | 2.14 ± 0.14 |
| Phenylalanine | 0.86 ± 0.03 | Alanine | 2.07 ± 0.18 |
| Tryptophan | 1.94 ± 0.15 | Proline | 4.32 ± 0.35 |
| Leucine | 1.83 ± 0.17 | Arginine | 2.69 ± 0.25 |
| Lysine | 2.25 ± 0.22 | Tyrosine | 0.95 ± 0.09 |
| Histidine | 5.35 ± 0.47 | Cystine | 1.10 ± 0.06 |
| **Total Essential AA** | 17.24 | **Total Non-Essential AA** | 21.52 |

The results are based on triplicate determinations ± SD.

### Fatty acid compositions

Table 4 summarizes 22 fatty acids from three different categories, namely saturated (SFA), monounsaturated (MUFA), and polyunsaturated (PUFA). Palmitic acid (C16:0) is the most abundant fatty acid in *Pantala* sp., which is consistent with data from the black soldier fly [40]. The second most abundant fatty acid is stearic acid (C18:0), followed by cis-9-oleic acid (C18:1n-9) and linolenic acid (C18:2n-6). These findings are similar to those reported previously in lesser mealworms (*Alphitobius diaperinus* Panzer; Coleoptera: Tenebrionidae) by Oonincx *et al*. [41]. In this insect, total SFA concentration was highest, followed by monounsaturated fatty acid (MUFA) and polyunsaturated fatty acid (PUFA). SFA concentrations ranged between 10.00 mg/100g and 1,300.00 mg/100g. Palmitic acid (C16:0) was the most abundant SFA in this insect, followed by stearic acid (C18:0) and heptadecanoic acid (C17:0). The MUFA concentration ranged between 10.00 mg/100g and 610.00 mg/100g. Cis-9-oleic acid (C18:1n-9c) was the most abundant MUFA in this insect, followed by palmitoleic acid (C16:1) and cis-10-Heptadecenoic acid (C17:1n-10). The PUFA content ranged from 13.33 mg/100g to 363.33 mg/100g. Linoleic acid (LA, C18:2n-6c) was the most abundant PUFA in this insect, followed by alpha-linolenic acid (ALA, C18:3n-3) and arachidonic acid (AA, C20:4n-6).

**Table 4.** Fatty acid concentration (mg/100g DW) of *Pantala* sp.

| Name | Formula | Concentration (mg/100g of DM) |
| --- | --- | --- |
| Lauric acid | C12:0 | 16.67 ± 5.77 |
| Tridecanoic acid | C13:0 | 10.00 ± 0.00 |
| Myristic acid | C14:0 | 93.33 ± 5.77 |
| Pentadecanoic acid | C15:0 | 156.67 ± 5.77 |
| Palmitic acid | C16:0 | 1,300.00 ± 45.83 |
| Heptadecanoic acid | C17:0 | 266.67 ± 5.77 |
| Stearic acid | C18:0 | 796.67 ± 32.15 |
| Arachidic acid | C20:0 | 93.33 ± 5.77 |
| Heneicosanoic acid | C21:0 | 26.67 ± 11.55 |
| Behenic acid | C22:0 | 73.33 ± 5.77 |
| **Total SFA** |  | 2,833.33 |
| Palmitoleic acid | C16:1 | 260.00±0.00 |
| cis-10-Heptadecenoic acid | C17:1n-10 | 144.33±5.77 |
| trans-9-Elaidic acid | C18:1n-9t | 13.33±5.77 |
| cis-9-Oleic acid | C18:1n-9c | 610.00±10.00 |
| cis-11-Eicosenoic acid | C20:1n-11 | 10.00±0.00 |
| **Total MUFAs** |  | 1,036.67 |
| Linoleic acid (LA) | C18:2n-6c (ω-6) | 363.33±32.15 |
| gamma-Linolenic acid (GLA) | C18:3n-6 (ω-6) | 23.33±5.77 |
| alpha-Linolenic acid (ALA) | C18:3n-3 (ω-3) | 176.67±20.82 |
| cis-11,14-Eicosadienoic acid | C20:2 | 20.00±0.00 |
| cis-8,11,14-Eicosatrienoic acid | C20:3n-6 (ω-6) | 13.33±5.77 |
| Arachidonic acid (AA) | C20:4n-6 (ω-6) | 123.33±15.28 |
| Eicosapentaenoic acid (EPA) | C20:5n-3 (ω-3) | 60.00±10.00 |
| **Omega 3 (n – 3)** |  | 236.67 |
| **Omega 6 (n – 6)** |  | 523.32 |
| **Total PUFAs** |  | 780.00 |
| **Total UFA** |  | 1,816.67 |
| **PUFA/SFA ratio** |  | 0.28 |

Results are in means of triplicate analysis ± SD (*n* =3). SFA (saturated fatty acid); MUFA (monounsaturated fatty acids); PUFA (polyunsaturated fatty acids); UFA (unsaturated fatty acid = MUFA + PUFA). The total omega-3 fatty acid composition was calculated as the ∑(alpha-linolenic acid [C18:3n3]+EPA [C20:5n3]). The total omega-6 fatty-acid composition was calculated as the ∑(linoleic acid [C18:2n6c]+ gamma-linolenic acid [C18:3n6]+cis-8,11,14-eicosatrienoic acid [C20:3n6]+arachidonic acid [C20:4n6]).

The current study demonstrates that edible aquatic insects have diverse fatty acid composition profiles. Omega-6 and omega-3 fatty acids are essential because humans, like all mammals, cannot produce them and must obtain them through diet. Linoleic acid (LA; 18:2n-6) represents omega-6 fatty acids, while alpha-linolenic acid (ALA; 18:3n-3) represents omega-3 fatty acids [42]. Among the detected n-6 PUFAs, LA (C18:2n-6c) was the most abundant. GLA (18:3n-6), AA (C20:4n-6), and eicosatrienoic acid are some other n-6 fatty acids (C20:3n-6). Linoleic acid was discovered in this species at a concentration of 363.33 mg/100g, which is much higher than that found in a commonly edible insect such as crickets (*Acheta domesticus* Linnaeus) [40]. These LA fatty acids are extremely important to humans because LA is the precursor of AA (C20:4n-6), which is the precursor of prostaglandin E2 (PGE2), which is also known as eicosanoids and is involved in the regulation of gene expression. Linoleic acid is also a structural component of cell membranes and plays an important role in cell signaling. Furthermore, high PUFA n-6 family intake has been shown to lower plasma cholesterol concentrations (lower risk of coronary heart disease) [43]. Consuming n-3 PUFAs has been shown to reduce the risk of diabetes by decreasing glucose intolerance, preventing insulin resistance, decreasing thrombotic tendency, and lowering blood pressure [44]. Surprisingly, we discovered a significant amount of ALA (C18:3n-3) in our current study. In contrast, Bophimai and Siri [45], Raksakantong *et al*. [46], and Ghosh *et al*. [47] discovered trace amounts of ALA in six edible dung beetle species (20.3-96.2 mg/100g DM), eight Thai edible terricolous insect species (2.46-39.82 mg/100g), and four commercial edible insects (10-110 mg/100g DM). Long-chain PUFA were typically found in aquatic insects, particularly omega-3 fatty acids such as ALA and EPA, whereas ALA was found in terrestrial species [48]. Aquatic insects obtain long-chain PUFA from their larvae and algae diets or synthesize them using the D5 and D6 desaturase enzymes [49]. Alpha-Linolenic acid is primarily used as a precursor in the synthesis of EPA. EPA has been shown to be a precursor of prostaglandin E3 (PGE3) and to be beneficial to the cardiovascular system. However, the recommended polyunsaturated to saturated fatty acid ratio (P/S ratio) for humans should be greater than 0.4 in order to reduce the risks of cardiovascular disease, cancer, and asthma, among other diseases [50]. Although *Pantala* sp. had a lower P/S ratio of 0.4, the normal P/S ration of meat was also around 0.1.

The total PUFA concentration of this species is comparable to raw lean meats such as mutton (673 mg/100g), lamb (603 mg/100g), and beef (448 mg/100g), and much higher than that found in some edible terricolous insects such as the june beetle (516.73 mg/100g), termite (465.06 mg/100g), cicada (213.15 mg/100g), longan stink bug (420.84 mg/100g) and short tailed cricket (771.63 mg/100g) that consumed in Thailand [46, 51]. As a result, direct consumption of edible aquatic insects could significantly contribute to human healthy fat requirements.

### Bioaccumulation

Heavy metals such as Pb, Cd, and As were found in *Pantala* sp. during the study (Table 5). Narzari and Sarmah [18] previously estimated the nutritional aspects of *Sympetrum* sp. (Odonata: Libellulidae), but toxic heavy metals were missing. In our study, arsenic concentrations in *Pantala* sp. were relatively high, followed by Cd, and Pb had a limit of detection of 0.05 mg kg_-1_. This result was comparable to that of Aydoğan *et al*. [53]. They discovered that some aquatic insects (Coleoptera: Hydrophilidae) from Turkey’s Karasu River had high arsenic levels (0.2 - 14.4 mg kg_-1_). Similarly, Addo-Bediako and Malakane [54] reported As levels (range: 7.3 - 32.326 mg kg_-1_) in Gomphidae (Odonata) larvae from the Blyde River that were higher than our results.

**Table 5.**
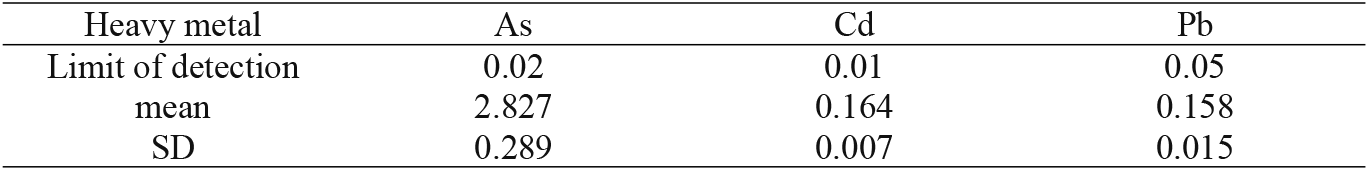
Mean ± SD (n = 3) of heavy metal concentrations detected in *Pantala* sp. (mg kg_-1_, dry weight) in the rice field.

Lead concentrations ranged from 0.142 to 0.172 mg kg_-1_, with an average of 0.158 ± 0.015 mg kg_-1_. Nasirian and Irvine [18] reported Pb and Cd levels in *Pantala flavescens* larvae (Odonata: Libellulidae) from the Shadegan and Hawr Al Azim wetlands that were similar to those in Table 5, though our Cd levels were higher. Furthermore, Choudhury *et al*. [5] discovered trace amounts of toxic heavy metals in the edible aquatic insect *Dytiscus marginalis* (Coleoptera: Dytiscidae). Local bioavailability, such as heavy metals from streams and sediments, could be one explanation. In particular, larger volumes of runoff due to rainfall and domestic waste enter the rice fields between May and October. Furthermore, these metals are commonly found in fertilizers and pesticides that are approved for use in Thailand. They could be present in the environment as natural or industrial contaminants [55]. Metal concentrations (i.e., Cd, Ni, Cr, As, Pb, Cu, Ti, Zn, and Mn) in aquatic insects may vary due to body size, life cycle stages, and different bioaccumulation patterns [56]. Thailand has no maximum levels for heavy metals in insect products. In contrast, the accumulation of Pb, Cd, and As in the edible aquatic insect *Pantala* sp. was compared to that of seafood. An edible aquatic insect from this study had a lower As content when compared to 13 different types of seafood (0.401 – 7.032 mg kg_-1_) and a lower Cd content than the bivalve group (blood cockle = 0.731 mg kg_-1_) from Rayong province, Thailand [57]. According to Thailand’s Ministry of Public Health [58], the Cd content in this sample did not exceed the 1.0 mg kg_-1_ maximum level for Cd in fish or the 2.0 mg kg_-1_ maximum level for cephalopods, marine bivalve mollusks, and gastropods. The highest level of Pb found in this insect was less than 0.3 mg kg_-1_, which is the maximum level for Pb in fish. Analyzing samples for arsenic content revealed that levels of 2.827 ± 0.289 mg kg_-1_ exceeded the permissible limits (2.0 mg kg_-1_) for food in Thailand. Arsenic is a common metalloid that can be found in low concentrations in rocks, soil, and natural water. Currently, no maximum levels of arsenic in food have been established in the EU [59]. The source of arsenic contamination was unknown, but it was thought to be a combination of fertilizers, pesticides, and industrial effluent [60]. These findings provided information about contaminated heavy metals in edible aquatic insects that could be used to improve food safety in this region. However, this paper does not indicate that the levels of toxic metals detected in edible aquatic insects pose a risk to human health, but they can cause bioaccumulation in consumers. Long-term accumulation of Cd in the human body can cause carcinogenesis in the skeletal and respiratory systems, as well as damage to the lungs and kidneys, whereas Pb can harm the gastrointestinal, neurologic, hematologic, cardiovascular, and renal systems. As can cause cancer in the skin, lungs, liver, and bladder, and it is toxic to the nervous and pulmonary systems [61].

## Conclusion

Eating insects has both benefits and disadvantages. According to this study, the edible aquatic insect found in rice fields contained high levels of protein, phosphorus, calcium, and iron. Furthermore, the tested insects act as reservoirs for essential amino acids (particularly lysine and histidine) and essential fatty acids (especially omega-3 and omega-6). Although insects have high nutritional value, edible insects must also be considered in terms of toxic metal bioaccumulation. This is about the possibility of food products entering the food chain. The concentration of heavy metals (particularly arsenic) in aquatic insects is determined by the metal concentration in the substrate in which they live and the stage of development of the insect. As a result, edible aquatic insects must assess their toxic elements as well, ensuring that the consumption of selected insects is safe.

## Acknowledgement

This research project is supported by the National Research Council of Thailand (NRCT): NRCT5-RGJ63002-041.

## References

1. FAO (Food and Agriculture Organization). Food energy-methods of analysis and conversion factors. 54: FAO Food and Nutrition paper 77. Rome, Italy. 2003.

2. Kim TK, Yong HI, Kim YB, Kim HW, Choi YS. Edible insects as a protein source: a review of public perception, processing technology, and research trends. Food science of animal resources. 2019; 39(4): 521–540.

3. Williams DD, Williams SS. Aquatic insects and their potential to contribute to the diet of the globally expanding human population. Insects. 2017; 8(3): 72.

4. Xiaoming C, Ying F, Hong Z. Review of the nutritive value of edible insects. In Forest Insects as Food: Humans Bite Back, Proceedings of a Workshop on Asia-Pacific Resources and Their Potential for Development; Durst, P.B., Johnson, D.V., Leslie, R.L., Shono, K., Eds.; FAO Regional Office for Asia and the Pacific: Bangkok, Thailand, 2010; pp. 85–92.

5. Choudhury K, Sarma D, Sapruna PJ, Soren AD. Proximate and mineral compositions of Samia cynthia ricini and Dytiscus marginalis, commonly consumed by the Bodo tribe in Assam, India. Bulletin of the National Research Centre. 2020; 44(1): 1–7.

6. Das JK, Hazarika AK. Macronutrient and mineral content of edible Coleopteran with reference to the Baksa district, India. The Clarion-International Multidisciplinary Journal. 2019; 8(2): 15–20.

7. Macadam CR, Stockan JA. The diversity of aquatic insects used as human food. Journal of Insects as Food and Feed. 2017; 3(3): 203–209.

8. Yhoung-Aree J, Viwatpanich K. Edible insects in the Lao PDR, Myanmar, Thailand and Vietnam. In M.G. Paoletti, ed. Ecological implications of mini–livestock. Potential of insects, rodents, frogs and snails, 2005; pp. 415–440. Enfield, New Hampshire, Science Publisher, Inc.

9. Hanboonsong Y. Edible insects and associated food habits in Thailand. In: Durst, P.B., Johnson, D.V., Leslie, R.N.and Shono, K. (eds.) Forest insects as food: humans bite back. Proceedings of a workshop on Asia-Pacific resources and their potential for development, 19-21 February 2008, Chiang Mai, Thailand. RAP Publication 2010/02. Food and Agriculture Organization of the United Nations, Bangkok, Thailand, pp. 173–182.

10. Ying F, Xiaoming C, Shaoyun W, Shoude Y, Yong C. Three edible Odonata species and their nutritive value. Forest Research. 2001; 14(4): 421–424.

11. Chakravorty J, Ghosh S, Meyer-Rochow VB. Comparative survey of entomophagy and entomotherapeutic practices in six tribes of Eastern Arunachal Pradesh (India). Journal of Ethnobiology and Ethnomedicine. 2013; 9(1): 1–12.

12. DeFoliart GR. Insects as human food: Gene DeFoliart discusses some nutritional and economic aspects. Crop protection. 1992; 11(5): 395–399.

13. Hanboonsong Y, Durst PB. Edible insects in Lao PDR: building on tradition to enhance food security. Food and Agriculture Organization of the United Nations, Bangkok, Thailand. 2014: 55 pp.

14. Barennes H, Phimmasane M, Rajaonarivo C. Insect consumption to address undernutrition, a national survey on the prevalence of insect consumption among adults and vendors in Laos. PloS one. 2015; 10(8): e0136458.

15. Sullivan DM, Carpenter DE. Methods of analysis for nutrition labeling chapter 6 p. 106. AOAC International, Arlington, Virginia USA. 1993.

16. Compendium of Methods for Food Analysis. 1st ed. Institute of Food Research and Product Development. Kasetsart University, Bangkok, Thailand. 2003.

17. Shantibala T, Lokeshwari RK, Debaraj H. Nutritional and antinutritional composition of the five species of aquatic edible insects consumed in Manipur, India. Journal of Insect Science. 2014; 14(1).

18. Narzari S, Sarmah J, Gupta P. Nutritional aspects of an aquatic edible insect Sympetrum sp. (Odonata: Libellulidae) of Assam, northeast India. International Journal of Food Sciences and Nutrition. 2017; 2(4): 38–42.

19. Narzari S, Sarmah J. Proximate composition of wild edible insects consumed by the Bodo tribe of Assam, India. International Journal of Bioassays. 2015; 4(7): 4050–4054.

20. Rahman A, Bordoloi S, Mazid S. Entomophagy practiced among the Tiwa community of Morigaon district, Assam. Journal of Entomology and Zoology Studies. 2018; 6(1): 484–486.

21. Alves AV, Sanjinez-Argandoña EJ, Linzmeier AM, Cardoso CAL, Macedo MLR. Food value of mealworm grown on Acrocomia aculeata pulp flour. PLoS One. 2016; 11(3): e0151275.

22. Thompson TE. Lipid. Encyclopedia Britannica. Available in: https://www.britannica.com/science/lipid. Accessed: 15/01/2021.

23. Akullo J, Agea JG, Obaa BB, Okwee-Acai J, Nakimbugwe D. Nutrient composition of commonly consumed edible insects in the Lango sub-region of northern Uganda. International Food Research Journal. 2018; 25(1): 159–166.

24. Payne CL, Scarborough P, Rayner M, Nonaka K. A systematic review of nutrient composition data available for twelve commercially available edible insects, and comparison with reference values. Trends in Food Science & Technology. 2016; 47: 69–77.

25. Adesina AJ. Proximate and anti-nutritional composition of two common edible insects: yam beetle (Heteroligus meles) and palm weevil (Rhynchophorus phoenicis). Elixir Food Science. 2012; 49: 9782–9786.

26. Idowu AB, Oliyide EO, Ademolu KO, Bamidele JA. Nutritional and anti-nutritional evaluation of three edible insects consumed by the Abeokuta community in Nigeria. International Journal of Tropical Insect Science. 2019; 39(2): 157–163.

27. Shrimanker I, Bhattarai S. Electrolytes. StatPearls [Internet]. Treasure Island (FL): StatPearls Publishing; 2021 Jan–. PMID: 31082167.

28. Ivanovitch K, Klaewkla J, Chongsuwat R, Viwatwongkasem C. Kitvorapat W. The intake of energy and selected nutrients by Thai urban sedentary workers: an evaluation of adherence to dietary recommendations. Journal of nutrition and metabolism. 2014.

29. Adeduntan SA. Nutritonal and Antinutritional Characteristics of Some Insects Foragaing in Akure Forest Reserve Ondo State, Nigeria. Journal of Food Technology. 2005; 3: 563–567.

30. Rumpold BA, Schlüter OK. Potential and challenges of insects as an innovative source for food and feed production. Innovative Food Science & Emerging Technologies. 2013; 17: 1–11.

31. Adámková A, Kouřimská L, Borkovcová M, Mlcek J, Bednárová M. Calcium in edible insects and its use in human nutrition. Potravinarstvo Slovak Journal of Food Sciences. 2014; 8(1): 233–238. https://doi.org/10.5219/366

32. Das JK, Hazarika AK. Quantitative Analysis of Mineral Content of Six Edible terrestrial Insects Commonly Consumed by ethnic people in Baksa District, Assam, India. Clarion: International Multidisciplinary Journal. 2017; 6(2).

33. Carter J, Peterson KE, Wiecha JL, Nobrega S, Gortmaker SL. Planet Health: an interdisciplinary curriculum for teaching middle school nutrition and physical activity. Human Kinetics. 2007.

34. Meyers, L. D., Hellwig, J. P., and Otten, J. J. (Eds.). 2006. Dietary reference intakes: the essential guide to nutrient requirements. National Academies Press.

35. National Research Council. Diet and health: implications for reducing chronic disease risk. National Academies Press. Available in: https://www.ncbi.nlm.nih.gov/books/NBK218735/. Accessed: 20/01/2021.

36. Mwangi MN, Oonincx DG, Stouten T, Veenenbos M, Melse-Boonstra A, Dicke M, Van Loon JJ. Insects as sources of iron and zinc in human nutrition. Nutrition research reviews. 2018; 31(2): 248–255.

37. Cameron M, Hofvander Y. Manual on feeding infants and young children. Second editions. FAO of the United Nations, Rome, Italy. 1980.

38. Igwe CU, Ujowundu CO, Nwaogu LA, Okwu GN. 2011. Chemical analysis of an edible African termite Macrotermes nigeriensis, a potential antidote to food security problem. Biochemistry and Analytical Biochemistry. 2011; 1(105): 2161–1009.

39. Longvah T, Mangthya K, Ramulu P. Nutrient composition and protein quality evaluation of eri silkworm (Samia ricinii) prepupae and pupae. Food Chemistry. 2011; 128(2): 400–403.

40. Bbosa T, Tamale Ndagire C, Muzira Mukisa I, Fiaboe KK, Nakimbugwe D. Nutritional characteristics of selected insects in Uganda for use as alternative protein sources in food and feed. Journal of Insect Science. 2019; 19(6): 23.

41. Oonincx DG, Laurent S, Veenenbos ME, van Loon JJ. Dietary enrichment of edible insects with omega 3 fatty acids. Insect Science. 2020; 27(3): 500–509.

42. Rossi R, Pastorelli G, Cannata S, Corino C. 2010. Recent advances in the use of fatty acids as supplements in pig diets: a review. Animal Feed Science and Technology. 2010; 162(1-2): 1–11.

43. Capra S. Nutrient reference values for Australia and New Zealand: Including recommended dietary intakes. 2006.

44. Ruxton CHS, Reed SC, Simpson MJA, Millington KJ. The health benefits of omega-3 polyunsaturated fatty acids: a review of the evidence. Journal of human nutrition and dietetics. 2004; 17(5): 449–459.

45. Bophimai P, Siri S. 2010. Fatty acid composition of some edible dung beetles in Thailand. International Food Research Journal. 2010; 17(4): 1025–1030.

46. Raksakantong P, Meeso N, Kubola J, Siriamornpun S. 2010. Fatty acids and proximate composition of eight Thai edible terricolous insects. Food Research International. 2010; 43(1): 350–355.

47. Ghosh S, Lee SM, Jung C, Meyer-Rochow VB. 2017. Nutritional composition of five commercial edible insects in South Korea. Journal of Asia-Pacific Entomology. 2017; 20(2): 686–694.

48. Twining CW, Brenna JT, Lawrence P, Shipley JR, Tollefson TN, Winkler DW. Omega-3 long-chain polyunsaturated fatty acids support aerial insectivore performance more than food quantity. Proceedings of the National Academy of Sciences. 2016; 113(39): 10920–10925.

49. Sprecher H. Metabolism of highly unsaturated n-3 and n-6 fatty acids. Biochem Biophys Acta. 2000; 1486: 219–231.

50. Milićević D, Vranić D, Mašić Z, Parunović N, Trbović D, Nedeljković-Trailović J, Petrović Z. The role of total fats, saturated/unsaturated fatty acids and cholesterol content in chicken meat as cardiovascular risk factors. Lipids in Health and Disease. 2014; 13(1): 1–12.

51. Williams P. Section 2: key nutrients delivered by red meat in the diet. Nutrition & Dietetics. 2007; 64(4): S113–S119.

52. Nasirian H, Irvine KN. Odonata larvae as a bioindicator of metal contamination in aquatic environments: application to ecologically important wetlands in Iran. Environmental monitoring and assessment. 2017; 189(9): 1–18.

53. Aydogan Z, Şişman T, İncekara Ü, Gürol A. Heavy metal accumulation in some aquatic insects (Coleoptera: Hydrophilidae) and tissues of Chondrostoma regium (Heckel, 1843) relevant to their concentration in water and sediments from Karasu River, Erzurum, Turkey. Environmental Science and Pollution Research. 2017; 24(10): 9566–9574.

54. Addo-Bediako A, Malakane K. Preliminary Assessment of Chemical Elements in Sediments and Larvae of Gomphidae (Odonata) from the Blyde River of the Olifants River System, South Africa. International Journal of Environmental Research and Public Health. 2020; 17(21): 8135.

55. Kolakowski BM, Johaniuk K, Zhang H, Yamamoto E. Analysis of Microbiological and Chemical Hazards in Edible Insects Available to Canadian Consumers. Journal of Food Protection. 2021.

56. Cid N, Ibáñez C, Palanques A, Prat N. Patterns of metal bioaccumulation in two filter-feeding macroinvertebrates: exposure distribution, inter-species differences and variability across developmental stages. Science of the Total Environment. 2010; 408(14): 2795–2806.

57. Kerdthep P, Tongyonk L, Rojanapantip L. Concentrations of cadmium and arsenic in seafood from Muang District, Rayong Province. Journal of Health Research. 2009; 23(4): 179–184.

58. Notification of Ministry of Public Health in Thailand. Standards for Contaminants in Food. Ministry of Public Health in Thailand. Available: http://www.food.fda.moph.go.th/law/data/announ_moph/V.English/P414_E.pdf. Accessed 01/08/2020.

59. European Food Safety Authority. Dietary exposure to inorganic arsenic in the European population. EFSA Journal. 2014; 12(3): 3597.

60. Choprathumma C, Thongkam T, Jaiyen C, Tusai T, Apilux A, Kladsomboon S. Determination of toxic heavy metal contaminated in food crops in Nakhon Pathom province, Thailand. Khon Kaen Agriculture Journal. 2019; 47 (1): 83–94.

61. Hung DQ, Nekrassova O, Compton RG. Analytical methods for inorganic arsenic in water: a review. Talanta. 2004; 64(2): 269–277.

